# The Generalized Data Model for Clinical Research

**DOI:** 10.1101/194597

**Authors:** Mark D. Danese, Marc Halperin, Jennifer Duryea, Ryan Duryea

## Abstract

1.

**1.1 Background:** Most healthcare data sources store information within their own unique schemas, making reliable and reproducible research challenging. Consequently, researchers have adopted various data models to improve the efficiency of research. Transforming and loading data into these models is a labor-intensive process that can alter the semantics of the original data. Therefore, we created a data model with a hierarchical structure that simplifies the transformation process and minimizes data alteration.

**1.2 Methods:** There were two design goals in constructing the tables and table relationships for the Generalized Data Model (GDM). The first was to focus on clinical codes in their original vocabularies to retain the original semantic representation of the data. The second was to retain hierarchical information present in the original data while retaining provenance. The model was tested by transforming synthetic Medicare data; Surveillance, Epidemiology, and End Results data linked to Medicare claims; and electronic health records from the Clinical Practice Research Datalink. We also tested a subsequent transformation from the GDM into the Sentinel data model.

**1.3 Results:** The resulting data model contains 19 tables, with the Clinical Codes, Contexts, and Collections tables serving as the core of the model, and containing most of the clinical, provenance, and hierarchical information. In addition, a Mapping table allows users to apply an arbitrarily complex set of relationships among vocabulary elements to facilitate automated analyses.

**1.4 Conclusions:** The GDM offers researchers a simpler process for transforming data, clear data provenance, and a path for users to transform their data into other data models. The GDM is designed to retain hierarchical relationships among data elements as well as the original semantic representation of the data, ensuring consistency in protocol implementation as part of a complete data pipeline for researchers.

## 2. Background

Healthcare data contains useful information for clinical researchers across a wide range of disciplines, including pharmacovigilance, epidemiology, and health services research. However, most data sources throughout the world store information within their own unique schemas, making it difficult to develop software tools that ensure reliable and reproducible research. One solution to this problem is to create data models that standardize the storage of both the data and the relationships among data elements.[1]

In healthcare, several commonly used data models include those supported by the following organizations: Informatics for Integrating Biology and the Bedside (i2b2) [2–4], Observational Health Data Sciences and Informatics (OHDSI, managing the OMOP [Observational Outcomes Medical Partnership] data model) [5–7], Sentinel [8–10], and PCORnet (Patient Centered Outcomes Research Network) [11, 12], among others. The first, and biggest, challenge with any data model is the process of migrating the raw (source) data into the data model, referred to as the “extract, transform, and load” (ETL) process. The ETL process is particularly burdensome when one has to support multiple, large data sources, and to update them regularly.[13]

Some aspects of transforming raw data into a particular data model are straight-forward, including reorganizing variables and standardizing their names. However, the most challenging aspect is standardizing the relationships among data elements without changing their meaning. Since different healthcare data sources encode relationships in different ways, the ETL process can lose information, or create inaccurate information. The best example is the process of creating a visit, a construct which, in most data models, is used to link information (e.g., diagnoses and procedures) on a per patient basis.

Visits are challenging because administrative claims allow facilities and practitioners to invoice separately for their portions of the same medical encounter, and allow practitioners to bill for multiple interactions on a single invoice.[14] Within the practitioner bills, individual procedures are linked to diagnosis codes, procedure modifiers, and costs. Consequently, a visit should link both the facility and the practitioner information without changing the existing practitioner-specified relationships between procedures, modifiers, diagnoses, and costs. Even electronic medical records can be challenging when each interaction with a different provider (e.g., nurse, physician, pharmacist, etc.) is recorded separately, requiring decisions to be made about defining a visit.

To minimize the need to encode specific relationships that may not exist in the source data, we created a data model with a hierarchical structure that minimizes changes to the meaning of the original data. This data model can serve both as a stand-alone data model for clinical researchers using observational data, as well as a storage model for later conversion into other data models.

## 3. Methods

In designing the Generalized Data Model (GDM) the primary use case was to allow clinical researchers using commonly available observational datasets to conduct research efficiently using a common framework. In particular, the GDM was designed to allow researchers to reuse an extensive, published body of existing algorithms for identifying clinical research constructs, including visits, that are expressed in the native vocabularies of the raw data. These algorithms require code sets, and may also require temporal logic (e.g., before, after, during, etc.), sequencing information (e.g., first, last, etc.), and provenance information (e.g., inpatient, outpatient, etc.). The GDM specifically considered both oncology research, which has its own specific vocabularies, and health services research. However, the model was designed so that these specific focus areas would not limit the design or use of the model.

### 3.1 Design goals

We initiated development of the GDM to make ETL specification and implementation easier for users who work with data models. There were two primary goals in defining the standard tables and table relationships for the GDM, described below.

#### 3.1.1 Focus on clinical codes in their original vocabularies

For clinical research, transparency and reproducibility are critically important. Therefore, the model is focused on the original (source) vocabularies to prevent the loss of the original semantic expression of the underlying clinical information. We also wanted all clinical codes (e.g., International Classification of Diseases [ICD], Current Procedural Terminology, National Drug Codes, etc.) to be easy to load into the data model and easy to query, because they represent the majority of electronic clinical information. Hence, the key organizing structure of the GDM is the placement of all clinical codes in a central “fact” table. This is not unlike the i2b2 data model that uses a fact table to store all “observations” from a source data set; however, the GDM was not designed as a star schema despite the similar idea of locating the most important data at the center of the data model.

We also considered interoperability as part of the design, but it was of secondary importance. Interoperability, like the construction of visits, requires establishing new connections (“mappings”) between the source vocabularies and a standard vocabulary such that a single query can operate across all data sources regardless of the source vocabulary. For international studies using different vocabularies, this might be a useful tool. However, given that every code isn’t yet mapped to a standard (e.g., OMOP has little in the way of procedure code mappings), and the maintenance required to support and update mappings, we designed the GDM to incorporate reliable cross-vocabulary mappings where they exist.

#### 3.1.2 Retain hierarchical information with provenance

The second goal was to capture important hierarchical relationships among data elements within a relational data structure. Based on the review of numerous data sources including Medicare, Surveillance Epidemiology and End Results (SEER) Medicare, Optum, Truven, JMDC (Japanese claims), and Clinical Practice Research Datalink (CPRD), we decided on a two-level hierarchy for grouping clinical codes, with the lower level table called Contexts and the higher-level table called Collections. This was based on common data structures where many related codes are recorded on a single record in the source data (Contexts table), and where these records are often grouped together (Collections table) based on clinical reporting or billing considerations. See Results for table definitions, and Figure 1 for a visual depiction of the hierarchical structure of the Contexts and Collections tables.

**Figure 1.**
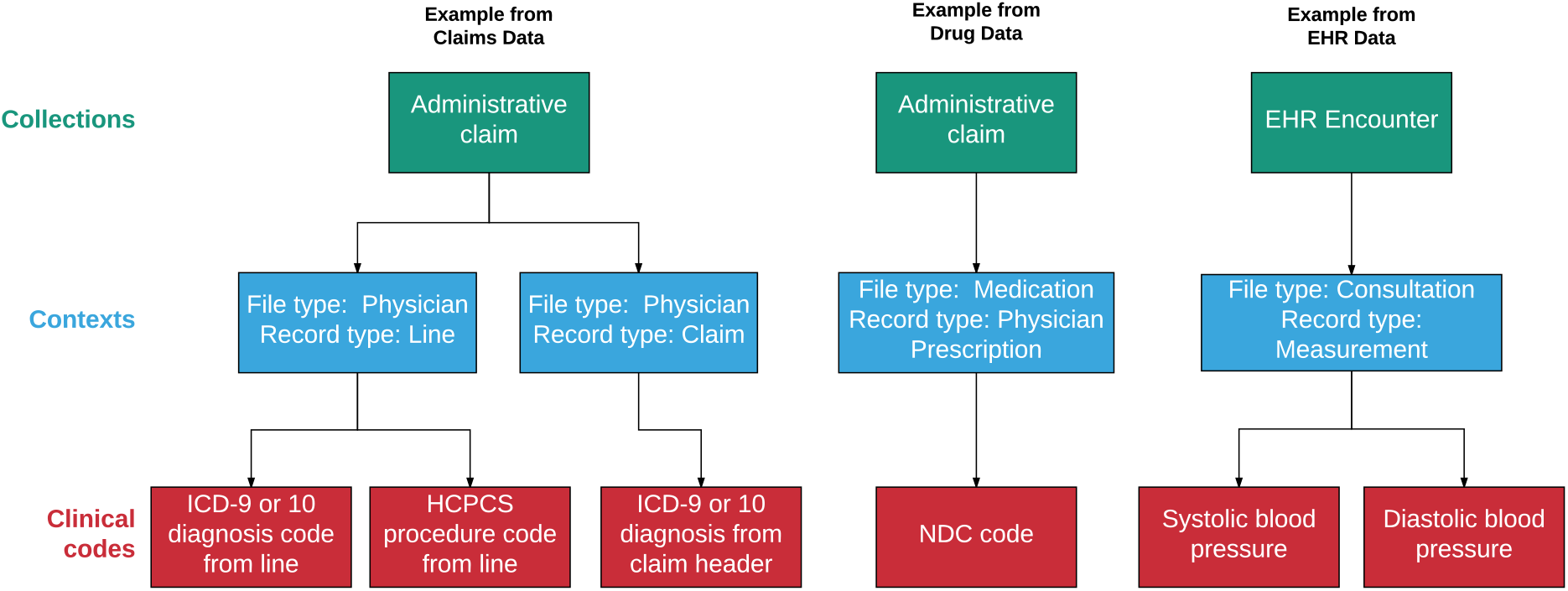
Relationships Among the Collections, Contexts, and Clinical Codes Tables. Note: EHR = electronic health record. HCPCS = Healthcare Common Procedure Coding System. NDC = National Drug Code. ICD = International Classification of Diseases. Figure does not contain specific data, but is intended to show the conceptual relationships among data elements across tables.

Our review of data sources suggested that the data model needed to support relatively few relationship types. The primary relationship represents data that is reported together or collected at the same time. One example of this includes a “line”, which occurs in claims data when one or more diagnosis codes, a procedure code, and a cost are all reported together. Another example includes laboratory values assessed at the same time (e.g., systolic and diastolic blood pressure) which could be considered to be co-reported. Also, a set of prescription refills could represent a linked set of records. Even records that contain pre-coordinated expressions (i.e., a linked set of codes used to provide clinical information akin to an English sentence) could also be stored in order by associating the codes with a single Context record.

We also included the provenance for each clinical code as part of Contexts, recording not only the type of relationship among elements within a Context as discussed above, but also the source file from which the data was abstracted. To minimize the loss of information when converting from the GDM to a data model that uses visits for organizing and consolidating most data relationships, the GDM does not require explicit visits (see Results). This is important because visits are not consistently defined among other data models, particularly for administrative claims data (see Discussion).

#### 3.1.3 Other Considerations

There are several other considerations made in building this data model, some of which were borrowed or adapted from other data models. For example, in addition to the cost table, we borrowed the OMOP idea to store all codes as “concept ids” (unique numeric identifiers for each code in each vocabulary to avoid conflicts between different vocabularies that use the same code). We also expanded upon the idea of OMOP “type_concept_ids” to track provenance within our data model. Finally, we allow flexibility in storing enrollment information in the Information Periods table using a “type_concept_id” so that the data can be used for different purposes (e.g., if a protocol does not require drug data, then enrollment in a drug plan should not be required). We also wanted to facilitate a straightforward, subsequent ETL process to other data models, including OMOP, Sentinel, and PCORnet.

We adapted the Payer Reimbursements table from the OMOP version 5.2 Cost table because it was the only data model with a cost table, and because we contributed substantially to its design. However, unlike the single OMOP cost table, we created two tables to accommodate both reimbursement-specific information, which has a well-defined structure, and all other kinds of economic information, which requires a very flexible structure. (The OMOP version 5.31 Cost table was redesigned to be more flexible, coincidentally resembling the GDM Costs table.)

### 3.2 Test Data

We tested the data model on three very different types of commonly available data used by clinical researchers: administrative claims data, EHR data, and cancer registry data. Claims in the United States are generally submitted electronically by the provider to the insurer using the American National Standards Institute (ANSI) 837P and 837I file specifications, which correspond to the CMS-1500 and UB04 paper forms.[15] Remittance information is sent from the insurer to the provider using the 835P and 835I specifications. However, actual claims data used for research is provided in a much simpler format. Based on experience developing and supporting software for submitting claims to insurers as well as creating ETL specifications for multiple commercial claims and EHR datasets using the OMOP data model, we determined that Medicare data is the most stringent test for transforming claims data because it contains the most information from the 837 and 835 files. For EHR data, we used the Clinical Practice Research Datalink (CPRD) data, because it is widely used for clinical research.[16] Finally, as part of our focus on oncology research, we included Surveillance, Epidemiology, and End Results (SEER) data[17] because SEER provides some of the most detailed cancer registry data available globally to clinical researchers which is challenging to incorporate into data models.

More specifically, we implemented a complete ETL process for the Medicare Synthetic Public Use Files (SynPUF). The SynPUF data are created from a 2.1-million-patient sample of Medicare beneficiaries from 2008 who were followed for three years, created to facilitate software development using Medicare data.[18, 19] We also implemented an ETL for SEER data linked to Medicare claims data [20] for 20,000 patients with small cell lung cancer, as part of an ongoing research project to describe patterns of care in that population. Finally, we developed a complete ETL for 140,000 CPRD patients for an ongoing research project evaluating outcomes associated with adherence to lipid-lowering medications. We also tested the feasibility of an ETL process to move SynPUF data from the GDM to the Sentinel data model (version 6.0) to ensure that the model did not contain any structural irregularities that would make it difficult to move data into other data model structures.

Finally, we conducted a test of information loss in the context of applying quality control to a study of mesothelioma patients. We conducted two analyses by separate people based on a written specification document using SEER Medicare data. The first was conducted using the source data and a combination of SAS and R code, and the second was conducted using the GDM version of the data and proprietary software. The analysis required the use of several SEER-specific fields, including the tumor sequence (first primary), histology, reporting type (microscopic confirmation), reporting source (not at death or autopsy), and tumor location data.

### 3.3 ETL Software

Our ETL process focused on the extraction of the source data and the transformation to the GDM data model, and saved tables as.csv files (i.e., it focused primarily on the E and T parts of the ETL). The ETL processes were built using R (version 3.4.4) and the data.table package (version 1.11.6).[21] R was selected because it is an open-source, cross-platform software package; because of its flexibility for composing ETL functions; and because of the availability of the data.table package as an in-memory database written in C for speed. The package itself is modular, and allows users to compose arbitrary ETL functions. Although the approach is different, the process is conceptually related to the dynamic ETL described by Ong, et al.[22]

## 4. Results

The resulting data model contains 19 tables (see hierarchical view in Figure 2). Details of the tables are provided in Supplementary Materials, and the most up-to-date version is available on a GitHub repository.[23] This repository will also contain links to any publicly available ETL specifications that we develop.

**Figure 2.**
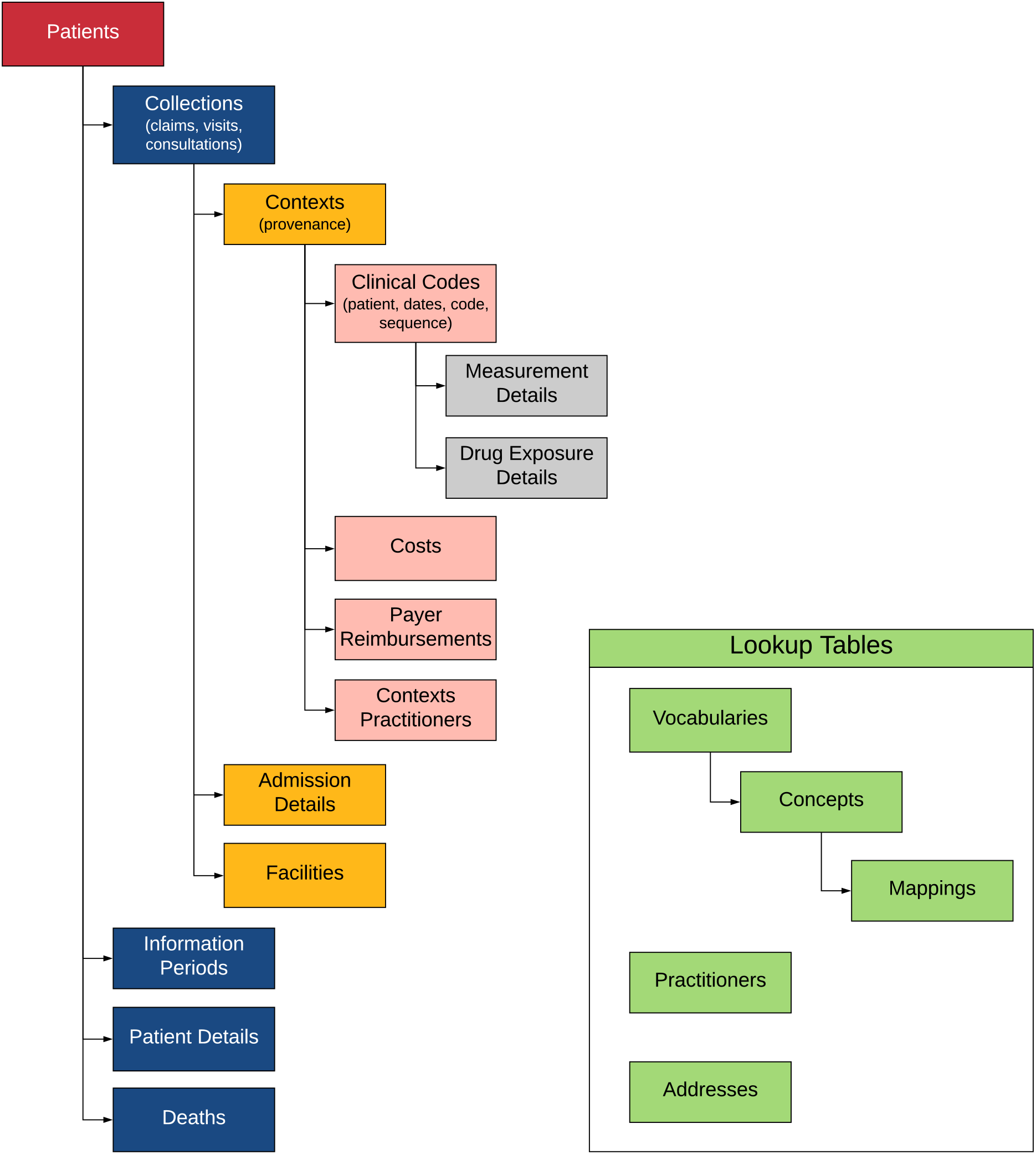
Hierarchical View of the Generalized Data Model. Note: Table names and key relationships among tables are depicted above. See Supplementary Materials for more detail on tables. Tables in green serve as lookup tables across the database. There is a single addresses table for unique addresses with relationships to Patients, Practitioners, and Facilities, and a single Practitioners table with relationships to Patients and Contexts Practitioners. The Contexts Practitioners table allows multiple practitioners to be associated with a Context record.

### 4.1 Tables

#### 4.1.1 Clinical Data

The Clinical Codes, Contexts, and Collections tables make up the core of the GDM (as shown in Figure 1). All clinical codes are stored in the Clinical Codes table. Each row of the Clinical Codes table contains a single code from the source data. In addition, each row also contains a patient id, the associated start and end dates for the record, a provenance concept id, and a sequence number. The sequence number allows codes to retain their order from the source data, as necessary. The most obvious example from billing data is diagnosis codes that are stored in numbered fields (e.g., diagnosis 1, diagnosis 2, etc.). But any set of ordered records could be stored this way, including groups of codes in a pre-coordinated expression. Grouping together ordered records in the Clinical Codes table is accomplished by associating them with the same id from the Contexts table. The provenance id allows for the specification of the type of record (e.g., admitting diagnosis, problem list diagnosis, etc.).

The Contexts table allows for grouping clinical codes and storing information about their origin. The record type concept id identifies the type of group that is stored. Examples might include lines from claims data where diagnoses, procedures, and other information are grouped, prescription and refill records that might be in electronic medical record or pharmacy data, or measurements of some kind from electronic health record or laboratory data (e.g., systolic and diastolic blood pressure, or a laboratory panel). In addition, the table stores the file name from the source data, the Center for Medicare and Medicaid Services place of service values (used for physician records since facility records to not have a place of service in claims data), and foreign keys to the care site and facility tables. The Contexts table also contains a patient id and both start and stop dates which could be different from the start and stop dates of the individual records from other tables to which the Contexts record is linked (e.g., a hospitalization may have different start and stop dates than the individual records within the hospitalization, as might occur with an in-hospital procedure performed on a single day of a multi-day hospitalization).

The Collections table represents a higher level of hierarchy for records in the Contexts table. That is, records in the Collections table represent groups of records from the Contexts table. This kind of grouping occurs when multiple billable units (“lines” or “details”) are combined into invoices (“claims”). It also occurs when prescriptions, laboratory measures, diagnoses and/or procedures are all recorded at a single office visit. In short, a Collection is typically a “claim” or a “visit” depending on whether the source data is administrative billing or electronic health record data. By using a hierarchical structure, the model avoids the requirement to construct "visits" from claims data which often leads to inaccuracy, loss of information, and complicated ETL processing. In the simplest possible case, it is possible to have a single record in the Clinical Codes table which is associated with a single Context record, which is associated with a single Collection record, as shown in Figure 1 for a drug record. The critical part of the ETL process, moving data into the Clinical Codes, Contexts, and Collections tables, is described in Figure 3 for the SynPUF data.

**Figure 3.**
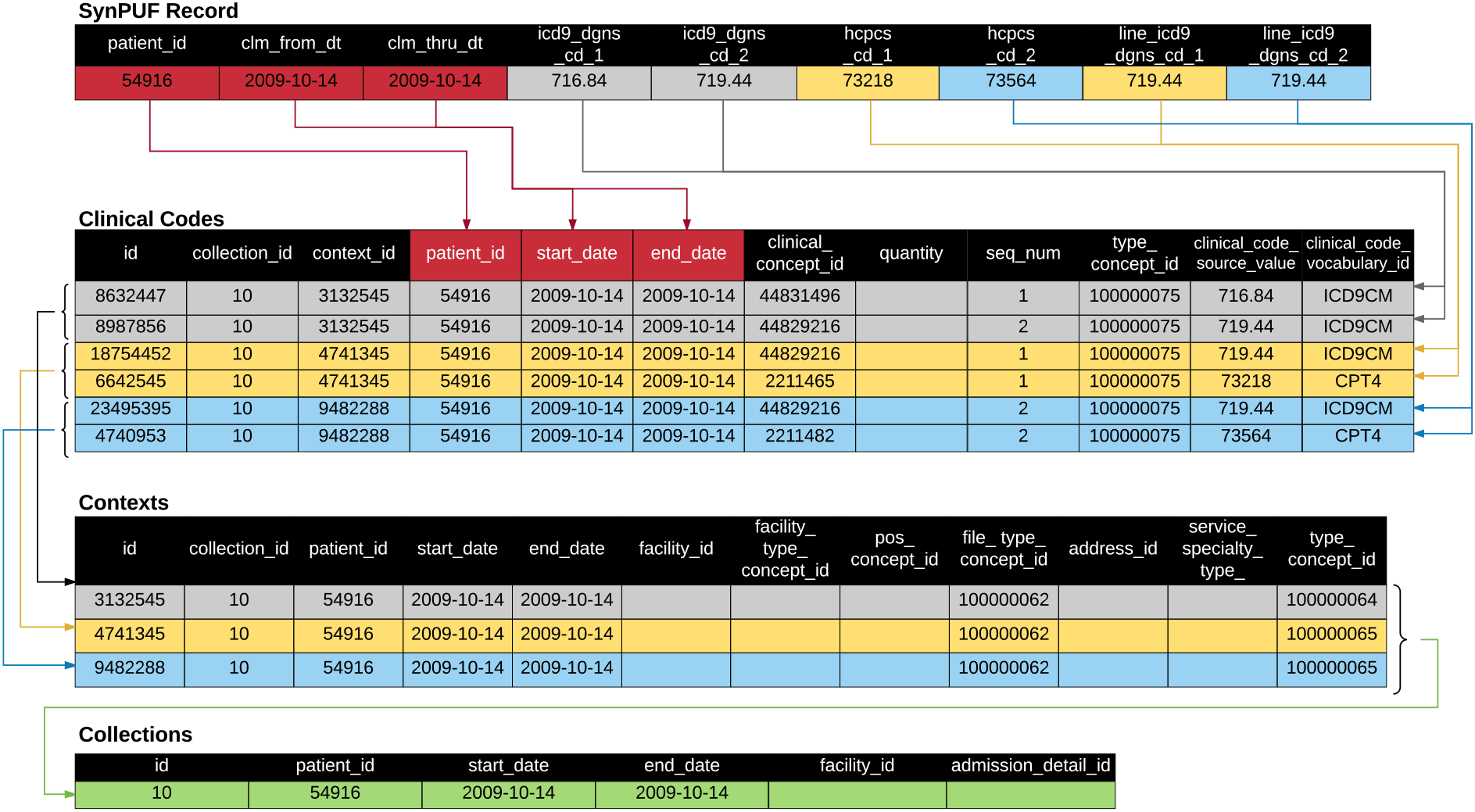
Visualization of the ETL Process for SynPUF Data. Note: Clinical codes are derived from a single row in the source data set (SynPUF record). Colored arrows indicate how each group of codes is used to create records. Each code from the original record gets its own row in the Clinical Codes table. Codes that are grouped together (e.g., line diagnosis 1 and procedure 1 in yellow) share the same context. In the Contexts table, type concept id ending in “64” indicates a claim level context, and the id ending in “65” indicates a line level context. The three contexts (groups of codes) share the same collection id.

The Details tables capture domain-specific information related to hospitalizations, drugs, and measurements. The Admissions Details table stores admissions and emergency department information that doesn’t fit in the Clinical Codes, Contexts, or Collections tables. It is designed to hold one admission per row. Each record in the Collections table for an inpatient admission links to this table. The Drug Exposure Details and Measurement Details contain information about medications and measurements (e.g., laboratory values). The Clinical Codes table contains foreign keys to these tables. We should also note that these two tables could be combined with the Clinical Codes table to make one larger table and improve query times on some database platforms. While this might require some minor modifications to the query, it wouldn’t change the underlying logic of the data model.

#### 4.1.2 Patient Data

The Patients table includes information about birth date, sex, race, ethnicity, address (via the Addresses table) and primary care provider (via Practitioners table). The Patient Details table allows a more flexible structure for timeless information like family history or simple genetic information. The Information Periods table captures periods of time during which the information in each table is relevant. This can include multiple records for each patient, including records for different enrollment types (e.g., Medicare Part A, Medicare Part B, or Medicare Managed Care) or this can be something as simple as a single date range of “up-to-standard” data as provided by the Clinical Practice Research Datalink. This table includes one row per patient for each unique combination of enrollment type and date range.

The Deaths table captures mortality information at the patient level, including date and cause(s) of death. This is typically populated from beneficiary or similar administrative data associated with the medical record. However, it is useful to check discharge status in the Admissions Details table as part of ETL process to ensure completeness. There are also diagnosis codes that indicate death. Deaths that are indicated by diagnosis codes should be in the Clinical Codes table and not be moved to the Deaths table. If needed, these codes can be identified using an appropriate algorithm (e.g., a set of ICD-9 codes, possibly with associated provenance specifications) to identify death as part of the identification of outcomes in an analysis.

There are two tables that store cost, charge, or payment data of some kind. The Payer Reimbursements table stores information from administrative claims data, with separate columns for each commonly used reimbursement element. All other financial information is stored in the Costs table, which is designed to support arbitrary cost types, and uses a “value_type_concept_id” to indicate the specific type. Costs may be present at a Context (line-item) or Collection (invoice) level. Therefore, this led us to align costs with the Contexts table. By evaluating the type of the context record, users can determine whether a cost is an aggregated construct or not. In administrative claims data, this means that each “line” (diagnosis and procedure) can have a cost record. For records that have costs only at the claim/header level (e.g., inpatient hospitalizations), only Contexts that refer to “claims” (i.e., a record_type_concept_id for “claim”) will have costs. For data with costs at both the line and claim/header level, costs can be distinguished by the Context type. In our experience, the sum of the line costs does not always equal the total cost, so depending on the research question, the researcher will need to determine whether claim, line, or both should be used. It is possible that each Clinical Code record sharing a single Contexts record could have a different cost; therefore, the two cost-related tables include a column to indicate the specific Clinical Code record to which the cost belongs. This might occur, for example, if multiple laboratory tests have different costs, but are share a common provenance (i.e., Contexts record).

#### 4.1.3 Facility and Practitioner Data

The Facilities table contains unique records for each facility where a patient is seen. The facility_type_concept_id should be used to describe the whole facility (e.g., Academic Medical Center or Community Medical Center). Specific departments in the facility should be entered in the Contexts table using the care_site_type_concept_id field. The Addresses table captures address information for practitioners and facilities, as well as patients.

The Contexts Practitioners table links one or more practitioners with a record in the Contexts table. Each record represents an encounter between a patient and a practitioner in a specific context. This role_type_concept_id in the table captures the role, if any, the practitioner played on the context (e.g., attending physician).

#### 4.1.4 Vocabulary Data

The Concepts table provides a unique numeric identifier (“concept_id”) for each source code in each vocabulary used in the data (see Table 1). Since queries against the GDM are intended to use the source codes, the Vocabulary table functions as a lookup table; therefore, the Concepts table does not have to be consistent across databases. However, there may be efficiencies in using a consistent set of identifiers for all entries from commonly used vocabularies. The specific vocabularies used in the data are provided in the Vocabularies table. The idea of having both Concepts and Vocabularies tables was adapted from the OMOP data models. As mentioned in Methods, the Mappings table allows for the expression of consistent concepts across databases.

**Table 1:** Example Concepts Table for Variables in the SEER Data.

| id | vocabulary_id | concept_code | concept_text |
| --- | --- | --- | --- |
| 100010943 | naaccr_grade | 1 | Grade I; grade i; grade 1; well differentiated; differentiated, NOS |
| 100010944 | naaccr_grade | 2 | Grade II; grade ii; grade 2; moderately differentiated; moderately well differentiated; intermediate differentiation |
| 100010945 | naaccr_grade | 3 | Grade III; grade iii; grade 3; poorly differentiated; differentiated |
| 100010946 | naaccr_grade | 4 | Grade IV; grade iv; grade 4; undifferentiated; anaplastic |
| 100010947 | naaccr_grade | 5 | T-cell; T-precursor |
| 100010948 | naaccr_grade | 6 | B-cell; Pre-B; B-Precursor |
| 100010949 | naaccr_grade | 7 | Null cell; Non T-non B; |
| 100010950 | naaccr_grade | 8 | N K cell (natural killer cell) |
| 100010951 | naaccr_grade | 9 | cell type not determined, not stated or not applicable |
| Note: SEER = Surveillance, Epidemiology and End Results, a collection of data from cancer registries across the United States. NAACCR = North American Association of Central Cancer Registries. |  |  |  |

The Mappings table is designed to express relationships among data elements. It can also be used to facilitate translation into other data models (see Table 2). In a few very simple cases like sex and race/ethnicity, we recommend concept mappings to a core set of values to make it easier for users of a protocol implementation software to filter patients by age, gender, and race/ethnicity using a simpler representation of the underlying information. The Mappings table also permits an arbitrarily complex set of relationships, along the lines of the approach taken with the OMOP model and the use of standard concepts for all data elements. By using a Mappings table, we reduce the need to re-map and re-load the entire dataset when new mappings become available. Regardless of how the Mappings table is used, the GDM still retains the original codes from the raw dataset.

**Table 2:**
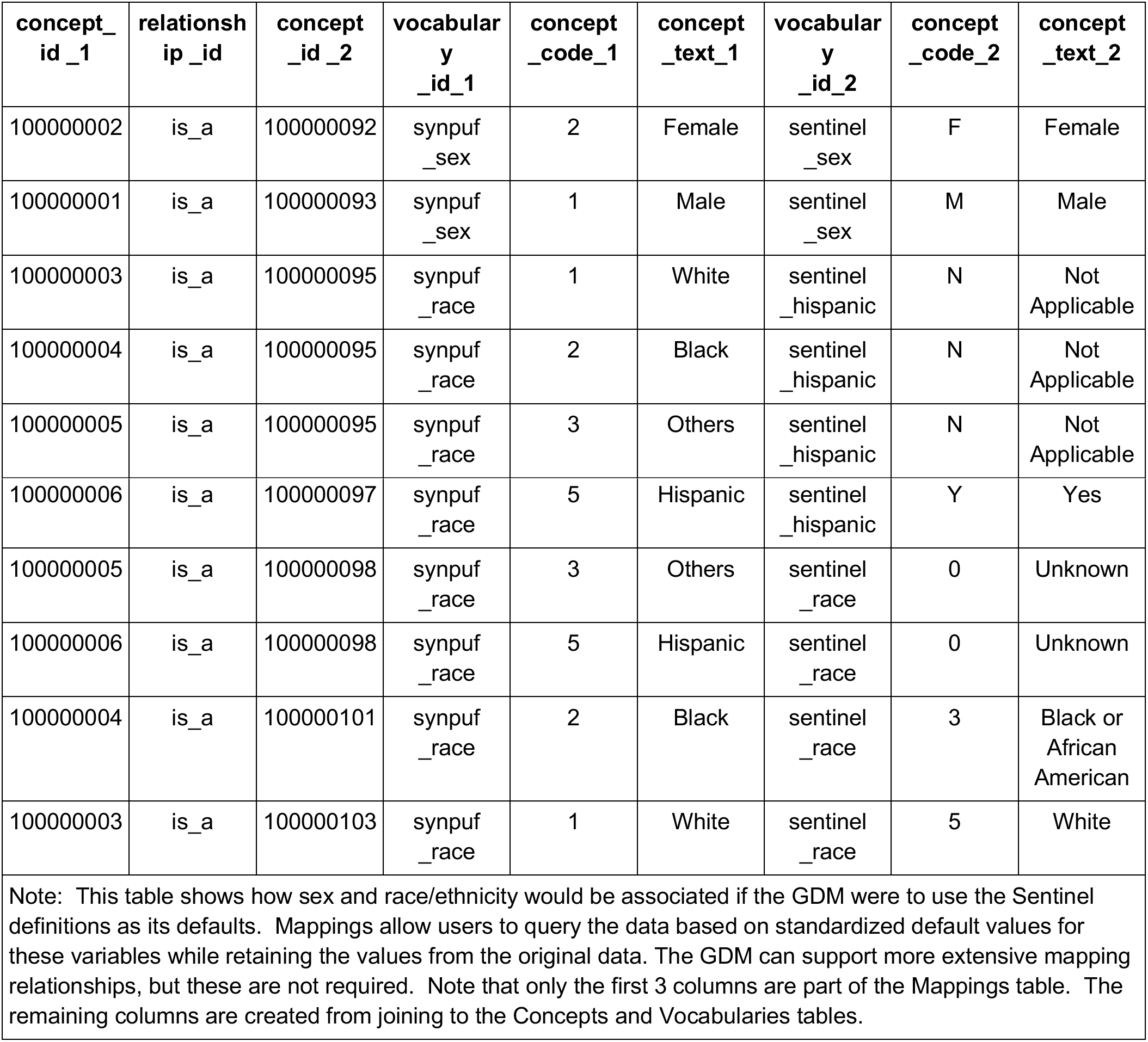
Example Mapping Table for Conversion to the Sentinel Model.

| concept_id_1 | relationship_id | concept_id_2 | vocabulary_id_1 | concept_code_1 | concept_text_1 | vocabulary_id_2 | concept_code_2 | concept_text_2 |
| --- | --- | --- | --- | --- | --- | --- | --- | --- |
| 100000002 | is_a | 100000092 | synpuf_sex | 2 | Female | sentinel_sex | F | Female |
| 100000001 | is_a | 100000093 | synpuf_sex | 1 | Male | sentinel_sex | M | Male |
| 100000003 | is_a | 100000095 | synpuf_race | 1 | White | sentinel_hispanic | N | Not Applicable |
| 100000004 | is_a | 100000095 | synpuf_race | 2 | Black | sentinel_hispanic | N | Not Applicable |
| 100000005 | is_a | 100000095 | synpuf_race | 3 | Others | sentinel_hispanic | N | Not Applicable |
| 100000006 | is_a | 100000097 | synpuf<br>_race | 5 | Hispanic | sentinel<br>_hispanic | Y | Yes |
| 100000005 | is_a | 100000098 | synpuf<br>_race | 3 | Others | sentinel<br>_race | 0 | Unknown |
| 100000006 | is_a | 100000098 | synpuf<br>_race | 5 | Hispanic | sentinel<br>_race | 0 | Unknown |
| 100000004 | is_a | 100000101 | synpuf<br>_race | 2 | Black | sentinel<br>_race | 3 | Black or<br>African<br>American |
| 100000003 | is_a | 100000103 | synpuf<br>_race | 1 | White | sentinel<br>_race | 5 | White |

Note: This table shows how sex and race/ethnicity would be associated if the GDM were to use the Sentinel definitions as its defaults. Mappings allow users to query the data based on standardized default values for these variables while retaining the values from the original data. The GDM can support more extensive mapping relationships, but these are not required. Note that only the first 3 columns are part of the Mappings table. The remaining columns are created from joining to the Concepts and Vocabularies tables.

### 4.2 ETL Results

We loaded SynPUF data and SEER Medicare data into the GDM. After downloading the data to a local server, the process of migrating the SynPUF data with 2.1 million patients of data to the GDM took approximately 8 hours on a Windows server with 4 cores and 128 Gb of RAM and conventional hard drives (running two files at a time in parallel). Most of the time was spent loading files into RAM and writing files to disk since the process of ETL with the GDM is primarily about relocating data.

SEER Medicare data for SCLC included approximately 20,000 patients and took less than 1 hour. Selected SEER data was included in the ETL process ignoring recoded versions of existing variables or variables used for consistency of interpretation over time. The ETL process focused on 31 key variables including histology, location, behavior, grade, stage, surgery, radiation, urban/rural status, and poverty indicators. Each SEER variable was included as a new vocabulary in the Concepts table (see Table 1).

CPRD data included approximately 140,000 patients and took approximately 2 hours. For the Test file which contains laboratory values and related measurements, we used Read codes in the Clinical Codes table; however, one could add the “entity types” (numeric values for laboratory values and other clinical measurements and assessments) to the Clinical Codes table as well, with both the Read code and the entity type associated with the same Context record and the same Measurement Details record. We used the entity types for all records in the CPRD Additional Clinical Details table. In all cases, the Mapping table allows for alternative relationships to be added to the data.

### 4.3 Information Loss

After reconciling differences in interpretation and resolving coding errors, we identified the identical cohort of patients when using the source data compared to using the same data in the GDM.

### 4.4 ETL from the GDM to Sentinel

We conducted an exploratory transformation from the GDM to Sentinel to ensure that it was feasible. The process of moving the data was conducted as follows. The transformations from the GDM Patients, Deaths, and Information Periods tables to Sentinel’s Demographic, Death, and Enrollment tables required renaming variables and mapping a source data vocabulary to a Sentinel vocabulary (e.g., SynPUF sex coding to Sentinel sex coding). The Sentinel Diagnosis, Procedure, and Dispensing tables were populated by splitting the GDM Clinical Codes table by clinical_code_source_vocabulary (e.g., ICD-9 codes were moved to the Sentinel Diagnosis table).

Populating the Sentinel Encounter table required records to be rolled up into a visit. To do this, the Contexts table was transformed into a “pre-Encounter” table with an encounter identifier set to the Contexts table identifier, with a similar process used for the Sentinel Procedure and Diagnosis tables. The “pre-Encounter” table was created with all of the specified columns and correctly mapped data, but had not yet grouped the records into visits. We applied logic based primarily on provenance information in the Contexts table to roll-up records into visits, and we created a new identifier in the Encounter table. Finally, the Diagnosis and Procedure tables were updated with new Encounter table identifier.

The remaining processing from the GDM to Sentinel involved vocabulary transformation since Sentinel has specific ways of representing concepts like sex which, in the GDM, are based on the source (e.g., male = 1 and female = 2) using a unique concept id in the Vocabulary table. We created records in the Mappings table from the SynPUF concepts to the Sentinel concepts (Table 2) to accomplish all needed mappings. Our ETL process then used those mappings to insert the correctly transformed variables from the GDM into the Sentinel tables during the ETL.

## 5. Discussion

The GDM is designed to allow clinical researchers to identify the clinical, resource utilization, and cost constructs needed for a wide range of epidemiological and health services research areas without altering the data’s original semantics by creating visits or domains, or performing substantial vocabulary mapping. This provides flexibility for researchers to study not only clinical encounters like outpatient visits, hospitalizations, emergency room visits, and episodes of care, but also more basic constructs like conditions or medication use. Its main goal is to simplify the location of the most important information for creating analysis data sets, which has the benefit of making ETL easier. It does this by using a hierarchical structure instead of visits. It tracks the provenance of the original data elements to enhance the reproducibility of studies. It includes a table to store relationships among data elements for standardized analyses. And it allows for a subsequent ETL process to other data models to provide researchers access to the analytical tools and frameworks associated with those models.

Because other data models (e.g., OMOP, Sentinel, PCORnet, and i2b2) use visits to connect patient-related information within the data model, our emphasis on avoiding visits deserves comment. Visits are seldom required for clinical research, unless the enumeration of explicit visits is the research topic itself. However, for most research projects, protocols require retrieval of the dates of specific, clinically relevant codes, perhaps with provenance or temporal constraints. Satisfying these criteria does not require knowledge of a visit, per se. It is a research project in and of itself to define visits, and their definitions are specific to the health services research question being investigated.[14] For example, a study of “emergency department” visits would need to consider at least four options to define a visit.[24] Data models that pre-define visits do not allow such flexibility.

The challenges with visits can best be seen by inspecting the guidelines for creating visits from each data model. In the Sentinel version 6 data model[10], a visit is defined as a unique combination of patient, start date, provider and visit type. Visit types are defined as Ambulatory, Emergency Department, Inpatient Hospital, Non-acute Institutional, and Other. Furthermore, “Multiple visits to the same provider on the same day should be considered one visit and should include all diagnoses and procedures that were recorded during those visits. Visits to different providers on the same day, such as a physician appointment that leads to a hospitalization, should be considered multiple encounters.”

PCORnet version 4.1 is similar to Sentinel.[12] However, PCORnet allows more visit types compared to PCORnet version 3, OMOP, and Sentinel. It includes Emergency Department Admit to Inpatient Stay, Observation Stay, and Institutional Professional Consult.

In the OMOP version 5.31 data model, a visit is defined for each “visit to a healthcare facility.” According to the specifications[6], in any single day, there can be more than one visit. One visit may involve multiple providers, in which case the ETL must either specify how a single provider is selected or leave it null. One visit may involve multiple care sites, in which case the ETL must either specify how a single site is selected or leave it null. Visits must be given one of the following visit types: Inpatient Visit, Outpatient Visit, Emergency Room Visit, Long Term Care Visit and Combined ER and Inpatient Visit. OMOP added an optional Visit Detail table in version 5.3, recognizing the two-level hierarchy common in US claims data.[6]

For i2b2, the specifications state a visit “… can involve a patient directly, such as a visit to a doctor’s office, or it can involve the patient indirectly, as in when several tests are run on a tube of the patient’s blood. More than one observation can be made during a visit. All visits must have a start date / time associated with them, but they may or may not have an end date. The visit record also contains specifics about the location of the session, such as the hospital or clinic the session occurred and whether the patient was an inpatient or an outpatient at the time of the visit.” There are no specified visit types, and the data model allows for an “unlimited number of optional columns but their data types and coding systems are specific to the local implementation.”[4]

Clearly, each data model has different perspectives on the definition of a visit. Such ambiguity can lead to differences in how tables are created in the ETL process. As a result, inconsistencies within or across data models can lead to differences in results, as has already been demonstrated.[25, 26] Laboratory records could be visits as with i2b2, or could be associated with visits as with other data models. Similarly, prescription, refill, and pharmacy dispensing records could be considered visits, or associated with visits. And other information, like family history, might not require a visit at all. In short, the most important structural component of other data models cannot be accurately and consistently defined, which affects the consistency of analyses across the data models, and makes translation among data models problematic. This also undermines provenance since each data model might answer the question of “where did this record come from” using different visit types. However, we note that these are semantic considerations and not technical limitations for record retrieval. For example, the i2b2 query platform recently has been extended to permit querying of OMOP and PCORnet data.[28]

One important consideration in using data models is their stability. It can be labor-intensive to keep data updated, and if both the data and the data model are changing, maintenance may be prohibitively time-consuming.[13] One of our intentions is that the GDM should remain stable over time; therefore, we incorporated separate Vocabulary and the Mappings tables which can be updated without running the ETL from the beginning. Hence, the GDM may be a useful, harmonized approach for data providers, compared to their various proprietary solutions. This contrasts with the OMOP data model which requires re-running the ETL when the vocabulary and domain mappings are updated.

The value of domains is that they allow data users to identify the necessary clinical information to extract for analysis and they facilitate interoperability. However, moving raw healthcare data into domains requires either mapping the entire vocabulary into a single domain, or mapping each individual code into a single domain. Placing codes in domain-specific tables can be particularly challenging when vocabularies cross domains (e.g., Read) or when individual codes are ambiguous (e.g., family history information). The GDM does not require domains or vocabulary mappings to be fully functional. The GDM only requires that users assign a unique number (concept id) to all unique source codes in a given dataset to ensure consistency in the data type for the codes. The vocabulary table is simply a look-up table for the codes and concept ids. Because of this, all codes in all vocabularies (e.g., ICD-9, HCPCS, etc.) in the source data will be retained unless there is an explicit decision to exclude a code. However, if needed, the GDM could support domains as an additional field in the Vocabulary table.

It is important to clarify the role of analyses in the ecosystem of data models. Neither the GDM nor any other data model is designed to support direct analyses of any sophistication on the entire database (excluding summary analyses to characterize the entire dataset). The role of the data model is to ease the extraction and organization of analysis data sets to address specific clinical research questions. The required analysis dataset structure depends on the specific analyses (e.g., prevalence, incidence, time to event, repeated measures, etc.) and is typically performed using R (OHDSI) or SAS (Sentinel). By starting with the GDM, researchers can develop tools to extract data directly, or implement the necessary transformations to migrate their data to other data models and make use of the tools for extraction and analysis offered by those models. While this requires another ETL process, or a database view to be created on the GDM, it facilitates access to existing analytical tools. Hence, the GDM can be used as a standardized waypoint in a data pipeline because the necessary information for other data models can be contained within the GDM as we found in our test of a GDM to Sentinel conversion.

We should also note that our approach to incorporating relationships into the data (i.e., our Mappings table) is not unique. Others have designed approaches that rely on semantic mappings to organize and extract data.[30] There are even methods to eliminate the need for both database reorganization and semantic mapping.[31] While these approaches may be more flexible and avoid cumbersome ETL and/or mapping processes, it is unclear how they fare with respect to the sensitivity and specificity of their exposure and outcome definitions making it challenging to understand or assess bias in their results.[32, 33]

Information loss and data quality assessment are challenging subjects. We designed the GDM to minimize information loss in the sense that any codes in the source data can be incorporated by creating entries in the Concepts, Vocabularies, and Clinical Codes tables. We also retained database specific provenance information by indicating the source file from which each data element is derived as well as the type of information that was derived. While we tested information loss in the context of a cohort study and found no problems, this is not a guarantee that all necessary information is, or can be, retained. A more robust assessment of data quality will be the subject of future research. However, our use of the SEER data is illustrative because detailed oncology data does not fit naturally into any of the other data models mentioned. Cancer registry data relies heavily on very specific vocabularies for location, histology, grade, staging, behavior, reporting source, microscopic confirmation and many other factors. Many of these don’t fit easily into the existing domain-based tables. The OMOP data model has a further complication in that the International Classification of Diseases for Oncology version 3 (ICD-O-3) which covers location, histology, grade, and behavior is not a standard vocabulary. Therefore, while the OMOP data model stores the concatenated source codes, work remains to be done to map all combinations to the proper standard vocabulary based on SNOMED. (This work is ongoing at the time of this writing.)

There are other limitations to the GDM. While we have tested it against data that is typically used by health services researchers and epidemiologists, there are likely to be specific data sets that will require modifications or improvements. The GDM does not yet include tables for patient reported outcomes, genomic data, or free text notes which are becoming more widely available for researchers. If other data models add support for these or other fields, this might require changes to the GDM to retain compatibility. For example, more detailed location information may need to be added for those with access to additional data (which is often limited due to privacy issues). While we have considered data from Japan and the United Kingdom, there are many data sources to which we did not have access that might require changes in the data model. Finally, while we have developed tools to extract analysis data sets from the GDM based on a protocol, they are not yet available publicly. (However, the ConceptQL language on which the tools are based is open-source.[34])

### 5.1 Conclusion

The GDM is designed to retain the relationships among data elements to the extent possible, facilitating ETL and protocol implementation as part of a complete data pipeline for clinical researchers using commonly available observational data. Furthermore, by avoiding the requirements to create visits and to use domains, it offers researchers a simpler process of standardizing the location of data in a defined structure and may make it easier for users to transform their data into other data models.

## 6. List of Abbreviations

**List of Abbreviations**

ANSI: American National Standards Institute
CMS: Centers for Medicare and Medicaid Services
CPRD: Clinical Practice Research Data link
EHR: Electronic Health Records
ETL: Extract Transform and Load
GDM: Generalized Data Model
ICD: International Classification of Diseases
i2b2: Informatics for Integrating Biology and the Bedside
NAACCR: American Association of Central Cancer Registries
OHDSI: Observational Health Data Science and Informatics
OMOP: Observational Outcomes Medical Partnership
PCORnet: Patient Centered Outcomes Research Network
SEER: Surveillance Epidemiology and End Results
SynPUF: Synthetic Public Use Files

## 7. Declarations

### 7.1 Ethics approval and consent to participate

A study protocol, an institutional review board exemption determination (Quorum IRB exemption determination #31309), and a data use agreement were required to access SEER-Medicare data. CPRD receives ethics approval to supply patient data for all protocols. Because of this, and the fact that all data are de-identified, no IRB approval or exemption is required. Our study protocol to access the CPRD data was reviewed by the CPRD Independent Scientific Advisory Committee. Provision of the CPRD data required a data use agreement. All transformations of the raw data were completed as part of the process of creating analysis data sets for approved study protocols. No clinical data was analyzed or reported for this study.

### 7.2 Consent for publication

Not applicable.

### 7.3 Availability of data and materials

The data model is publicly available. The raw data is not available due to privacy reasons, except for the Medicare Synthetic Public Use data. See **Ethics approval and consent to participate** for details on SEER Medicare and CPRD data acquisition, and **References** for a specific hyperlink to the Synthetic Public Use data.

## 7.5 Acknowledgements

We gratefully acknowledge the influence of the open-source OMOP model specifications on our thinking in creating our data model. In addition, we acknowledge the influence of Sentinel, PCORnet, and i2b2 on our approach, although most of our data model was designed prior to reviewing these models in detail. We also thank Chris Adamson for helpful discussions about organizing the data model in different ways. At the time of writing, all references to the concepts table refer to the OMOP version 5.20 vocabulary table maintained by OHDSI. However, there is no reason that a user could not create their own system of codes with unique identifiers across vocabularies, or use the codes from the National Library of Medicine Metathesaurus.[35]

## 7.6 Authors’ information

MD is an epidemiologist who has worked with a wide variety of clinical data sources across therapeutic areas. JD and RD have extensive experience designing software for providers to submit medical bills to insurers. JD has constructed and/or substantially revised OMOP ETL specifications for many commercially available data sources. MD, JD, and RD are collaborators in the Observational Data Health Sciences and Informatics organization.

## 7.7 Authors’ contributions

MD, MH, RD, and JD contributed to the design of the data model. MH wrote the software code for data transformations. All authors have read and approved the manuscript.

