## Supplementary Materials for "The Generalized Data Model for Clinical Research"

Title: A Generalized Data Model to Facilitate Biomedical Research

### 1. Generalized Data Model Tables

#### Table 1: Clinical Codes

| Column | Type | Description | Foreign Key | Required |
|---|---|---|---|---|
| id | serial | Surrogate key for record | | x |
| collection_id | bigint | FK reference to collections table | collections | x |
| context_id | bigint | FK reference to contexts table | contexts | x |
| patient_id | bigint | FK reference to patients table | patients | x |
| start_date | date | Start date of record (yyyy-mm-dd) | | x |
| end_date | date | End date of record (yyyy-mm-dd) | | x |
| clinical_code_concept_id | bigint | FK reference to concepts table for the code assigned to the record | concepts | x |
| quantity | bigint | Quantity, if available (e.g., procedures) | | |
| seq_num | int | The sequence number for the variable assigned (e.g. dx3 gets sequence number 3) | | |
| provenance_concept_id | bigint | Additional type information (ex: primary, admitting, problem list, etc) | concepts | |
| clinical_code_source_value | text | Source code from raw data | | x |
| clinical_code_vocabulary_id | text | Vocabulary the clinical code comes from | vocabularies | x |
| measurement_detail_id | bigint | FK reference to measurement_details table | measurement_details | |
| drug_exposure_detail_id | bigint | FK reference to drug_exposure_details table | drug_exposure_details | |

- Stores clinical codes from all types of records including procedures, diagnoses, drugs, laboratory records and other sources. Some common vocabularies include ICD-9, ICD-10, SNOMED, Read, HCPCS, CPT, NDC, and LOINC
- Ignores semantic distinctions about the type of information represented within a vocabulary because most vocabularies contain information from more than one domain
- One record generated for each individual code in the raw data

- Extra detail can be found about a code in the measurement_details and drug_exposure_details tables if that information exists

**Table 2: Contexts**

| Column | Type | Description | Foreign Key | Required |
|---|---|---|---|---|
| id | serial | Surrogate key for record | | x |
| collection_id | bigint | FK reference to collections table | collections | x |
| patient_id | bigint | FK to reference to patients table | patients | x |
| start_date | date | Start date of record (yyyy-mm-dd) | | x |
| end_date | date | End date of record (yyyy-mm-dd) | | x |
| facility_id | bigint | FK reference to facilities table | facilities | |
| care_site_type_concept_id | bigint | FK reference to concepts table representing the care site type within the facility | concepts | |
| pos_concept_id | bigint | FK reference to concepts table representing the place of service associated with this record | concepts | |
| source_type_concept_id | bigint | FK reference to concepts table representing the file name (e.g MEDPAR) and concatenate the subset of the file used to result in MEDPAR_SNF | concepts | x |
| service_specialty_type_concept_id | bigint | FK reference to concepts table representing the specialty type for the services/diagnoses associated with this record | concepts | |
| record_type_concept_id | bigint | FK reference to concepts table representing the type of contexts the record represents (line, claim, etc.) | concepts | x |

- Stores information about the context of the clinical_codes and costs
- Used to group clinical_codes typically occurring on the same day or at the same time (e.g., a diagnosis and a procedure, or a systolic and diastolic blood pressure)
- contexts records are always linked to a collection records
- care_site_type_concept_id is used to describe the department in which the service was performed

**Table 3: Collections**

| Column | Type | Description | Foreign Key | Required |
|---|---|---|---|---|
| id | serial | Surrogate key for record | | x |
| patient_id | bigint | FK to reference to patients table | patients | x |
| start_date | date | Start date of record (yyyy-mm-dd) | | x |
| end_date | date | End date of record (yyyy-mm-dd) | | x |
| duration | float | Duration of collection. (e.g. hospitalization length of stay) | | |
| duration_unit_concept_id | bigint | FK reference to concepts table for the unit of duration | concepts | |
| facility_id | bigint | FK reference to facilities table | facilities | |
| admission_detail_id | bigint | FK reference to admission_details table | admission_details | |
| collection_type_concept_id | bigint | FK reference to concepts table representing the type of collection this record represents | concepts | |

- Used to group contexts records
- For claims, records the claim level information (also referred to as "headers" in some databases)
    - Use claim from and thru date for start and end date, if available
    - Admit and discharge dates should go in the admission_details table unless those are the only dates for the records in which case they should be entered into both the collections and admission_details tables
- For EHR, records the visit level information

**Table 4: Measurement Details**

| Column | Type | Description | Foreign Key | Required |
|---|---|---|---|---|
| id | serial | Surrogate key for record | | x |
| patient_id | bigint | FK reference to patients table | patients | x |
| result_as_number | float | The observation result stored as a number, applicable to observations where the result is expressed as a numeric value | | |
| result_as_string | text | The observation result stored as a string, applicable to observations where the result is expressed as verbatim text | | |
| result_as_concept_id | bigint | FK reference to concepts table for the result associated with the detail_concept_id (e.g., positive/negative, present/absent, low/high, etc.) | concepts | |
| result_modifier_concept_id | bigint | FK reference to concepts table for result modifier (=, <, >, etc.) | concepts | |
| unit_concept_id | bigint | FK reference to concepts table for the measurement units (e.g., mmol/L, mg/dL, etc.) | concepts | |
| normal_range_low | float | Lower bound of the normal reference range assigned by the laboratory | | |
| normal_range_high | float | Upper bound of the normal reference range assigned by the laboratory | | |
| normal_range_low_modifier_concept_id | bigint | FK reference to concepts table for result modifier (=, <, >, etc.) | concepts | |
| normal_range_high_modifier_concept_id | bigint | FK reference to concepts table for result modifier (=, <, >, etc.) | concepts | |

- Stores additional information related to measurements, observations, status, and specifications
- Text-based vocabularies are sufficient, but could also be mapped to LOINC and stored in the mappings table (e.g., laboratory data indexed by text names for the lab results)
- Other vocabularies should be included in their original system (e.g., oncology may be comprised of separate vocabularies for location, histology, grade, behavior, etc.)

- This could be implemented by making variable names a vocabulary in themselves, depending on the use case

**Table 5: Drug Exposure Details**

| Column | Type | Description | Foreign Key | Required |
|---|---|---|---|---|
| id | serial | Surrogate key for record | | x |
| patient_id | bigint | FK to reference to patients table | patients | x |
| refills | int | The number of refills after the initial prescription; the initial prescription is not counted (i.e., values start with 0) | | |
| days_supply | int | The number of days of supply as recorded in the original prescription or dispensing record | | |
| number_per_day | float | The number of pills taken per day | | |
| dose_form_concept_id | bigint | FK reference to concepts table for the form of the drug (capsule, injection, etc.) | concepts | |
| dose_unit_concept_id | bigint | FK reference to concepts table for the units in which the dose_value is expressed | concepts | |
| route_concept_id | bigint | FK reference to concepts table for route in which drug is given | concepts | |
| dose_value | float | Numeric value for the dose of the drug | | |
| strength_source_value | text | Drug strength as reported in the raw data. This can include both dose value and units | | |
| ingredient_source_value | text | Ingredient/Generic name of drug as reported in the raw data | | |
| drug_name_source_value | text | Product/Brand name of drug as reported in the raw data | | |

- Designed to capture extra details about drug-specific clinical_codes
- The quantity of a drug is stored in the clinical_codes quantity field

**Table 6: Admission Details**

| Column | Type | Description | Foreign Key | Required |
|---|---|---|---|---|
| id | serial | Surrogate key for record | | x |
| patient_id | bigint | FK reference to patients table | patients | x |
| admission_date | date | Date of admission (yyyy-mm-dd) | | x |
| discharge_date | date | Date of discharge (yyyy-mm-dd) | | x |
| admit_source_concept_id | bigint | Database specific code indicating source of admission (e.g., ER visit, transfer, etc.) | | |
| discharge_location_concept_id | bigint | Database specific code indicating discharge location (e.g., death, home, transfer, long-term care, etc.) | | |
| admission_type_concept_id | bigint | FK reference to concepts table representing the type of admission the record is (Emergency, Elective, etc.) | concepts | |

- Captures details about admissions and emergency department encounters that cannot be stored in the clinical_codes, contexts, or collections tables
- One row per admission
- Each admission record in the collections table will link to this table

**Table 7: Payer Reimbursements**

| Column | Type | Description | Foreign Key | Required |
|---|---|---|---|---|
| id | serial | A unique identifier for each COST record | | x |
| context_id | bigint | FK reference to context table | contexts | x |
| patient_id | bigint | FK to reference to patients table | patients | x |
| clinical_code_id | bigint | FK reference to clinical_codes table to be used if a specific code is the direct cause for the reimbursement | clinical_codes | |
| currency_concept_id | bigint | FK reference to concepts table for the 3-letter code used to delineate international currencies (e.g., USD = US Dollar) | concepts | x |
| total_charged | float | The total amount charged by the provider of the good/service (e.g. hospital, physician pharmacy, dme provider) billed to a payer. | | |
| total_paid | float | The total amount paid from all payers for the expenses of the service/device/drug. | | |
| paid_by_payer | float | The amount paid by the Payer for the service/device/drug. | | |
| paid_by_patient | float | The total amount paid by the patient as a share of the expenses. | | |
| paid_patient_copay | float | The amount paid by the patient as a fixed contribution to the expenses. | | |
| paid_patient_coinsurance | float | The amount paid by the patient as a joint assumption of risk. | | |
| paid_patient_deductible | float | The amount paid by the patient that is counted toward the deductible defined by the Payer Plan. | | |
| paid_by_primary | float | The amount paid by a primary Payer through the coordination of benefits. | | |

| Column | Type | Description | Foreign Key | Required |
|---|---|---|---|---|
| paid_ingredient_cost | float | The amount paid by the Payer to a pharmacy for the drug, excluding the amount paid for dispensing the drug. | | |
| paid_dispensing_fee | float | The amount paid by the Payer to a pharmacy for dispensing a drug, excluding the amount paid for the drug ingredient. | | |
| information_period_id | float | FK reference to the information_periods table | | |
| amount_allowed | float | The contracted amount agreed between the payer and provider. | | |

- To store reimbursed costs (i.e., charges, paid amounts, and/or costs) for each provided service
- All reimbursed costs are linked to a record in the contexts table which identifies the type of cost (generally a line-level or claim-level cost)
- Note that claim-level costs do not always sum to the individual line-level costs, so caution should be used when querying records

**Table 8: Costs**

| Column | Type | Description | Foreign Key | Required |
|---|---|---|---|---|
| id | serial | A unique identifier for each COST record | | x |
| context_id | bigint | FK reference to context table | contexts | x |
| patient_id | bigint | FK to reference to patients table | patients | x |
| clinical_code_id | bigint | FK reference to clinical_codes table to be used if a specific code is the direct cause for the reimbursement | clinical_codes | |
| currency_concept_id | bigint | FK reference to concepts table for the 3-letter code used to delineate international currencies (e.g., USD = US Dollar) | concepts | x |
| cost_base | text | Defines the basis for the cost in the table (e.g., 2013 for a specific cost-to-charge ratio, or a specific cost from an external cost | | |
| value | float | Cost value | | x |
| value_type_concept_id | bigint | FK reference to concepts table to concept that defines the type of economic information in the value field (e.g., cost-to-charge ratio, calculated cost, reported cost) | concepts | x |

- Used to capture all non-reimbursement costs
- Example of things captured in this table are things like cost-to-charge ratio, calculated cost (for situations where the ETL process calculates a cost based on the available data), reported cost (where the ETL process imputes a cost from another source), and some other things that may become apparent with more use cases.

**Table 9: Patients**

| Column | Type | Description | Foreign Key | Required |
|---|---|---|---|---|
| id | serial | A unique identifier for each patient | | x |
| gender_concept_id | bigint | A foreign key that refers to an identifier in the concepts table for the unique gender of the patient | concepts | |
| birth_date | date | Date of birth (yyyy-mm-dd) | | |
| race_concept_id | bigint | A foreign key that refers to an identifier in the concepts table for the unique race of the patient | concepts | |
| ethnicity_concept_id | bigint | A foreign key that refers to an identifier in the concepts table for the ethnicity of the patient | concepts | |
| address_id | bigint | A foreign key to the place of residency for the patient in the location table, where the detailed address information is stored | addresses | |
| practitioner_id | bigint | A foreign key to the primary care practitioner the patient is seeing in<br>the practitioners table | practitioners | |
| patient_id_source_value | text | Originial patient identifier defined in the source data | | x |

- Demographic information about the patients in the data
- The column for *practitioner_id* is intended for situations where there is a defined primary care practitioner (e.g., HMO or CPRD data)

**Table 10: Patient Details**

| Column | Type | Description | Foreign Key | Required |
|---|---|---|---|---|
| id | serial | A unique identifier for each patient_detail | | x |
| patient_id | bigint | FK reference to patients table | patients | x |
| patient_detail_concept_id | bigint | FK reference to concepts table for the code assigned to the record | concepts | x |
| patient_detail_source_value | text | Source code from raw data | | x |
| patient_detail_vocabulary_id | text | Vocabulary the patient detail comes from | vocabularies | x |

- Extra information about a patient that doesn't fit in the patients table

**Table 11: Information Periods**

| Column | Type | Description | Foreign Key | Required |
|---|---|---|---|---|
| id | serial | Surrogate key for record | | x |
| patient_id | bigint | FK reference to patients table | patients | x |
| start_date | date | Start date of record (yyyy-mm-dd) | | x |
| end_date | date | End date of record (yyyy-mm-dd) | | x |
| information_type_concept_id | bigint | FK reference to concepts table representing the information type (e.g., insurance coverage, hospital data, up-to-standard date) | concepts | x |

- Captures periods for which information in each table is relevant for each person
- Could include enrollment types (e.g., Part A, Part B, HMO) or just "observable" (as with up-to-standard data in CPRD)
- One row per patient per enrollment/information period type

**Table 12: Deaths**

| Column | Type | Description | Foreign Key | Required |
|---|---|---|---|---|
| id | serial | Surrogate key for record | | x |
| patient_id | bigint | FK reference to patients table | patients | x |
| date | date | Date of death (yyyy-mm-dd) | | x |
| cause_concept_id | bigint | FK reference to concepts table for cause of death (typically ICD-9 or ICD-10 code) | concepts | |
| cause_type_concept_id | bigint | FK reference to concepts table for the type of cause of death (e.g. primary, secondary, etc.) | concepts | |
| practitioner_id | bigint | FK reference to practitioners table | practitioners | |

- Stores mortality information including date of death and cause(s) of death
- Commonly populated from beneficiary or similar administrative data associated with the medical record
- Deaths identified from diagnosis codes or discharge status are not necessary since such records are in the clinical_codes and admission_details tables and can be queried separately

**Table 13: Contexts Practitioners**

| Column | Type | Description | Foreign Key | Required |
|---|---|---|---|---|
| context_id | bigint | FK reference to contexts table | contexts | x |
| practitioner_id | bigint | FK reference to practitioners table | practitioners | x |
| role_type_concept_id | text | Roles practitioners can play in an encounter | | |
| specialty_type_concept_id | bigint | FK reference to concepts table representing the practitioner's specialty type for the services/diagnoses associated with this record | concepts | |

- Links one or more practitioners with a contexts record
- Each record represents an encounter between a patient and a practitioner on a specific context
- Captures the role, if any, the practitioner played on the context (e.g., attending physician)

**Table 14: Practitioners**

| Column | Type | Description | Foreign Key | Required |
|---|---|---|---|---|
| id | serial | A unique identifier for each practitioner | | x |
| practitioner_name | text | practitioners name, if available | | |
| primary_identifier | text | Primary practitioner identifier | | x |
| primary_identifier_type | text | Type of identifier specified in primary identifier field (UPIN, NPI, etc) | | x |
| secondary_identifier | text | Secondary practitioner identifier (Optional) | | |
| secondary_identifier_type | text | Type of identifier specified in secondary identifier field (UPIN, NPI, etc) | | |
| specialty_concept_id | bigint | A foreign key to an identifier in the concepts table for specialty | concepts | |
| address_id | bigint | A foreign key to the address of the location where the practitioner is practicing | addresses | |
| birth_date | date | Date of birth (yyyy-mm-dd) | | |
| gender_concept_id | bigint | A foreign key that refers to an identifier in the concepts table for the unique gender of the person | concepts | |

- All non-facility practitioners (i.e., physicians, etc.) are listed

**Table 15: Addresses**

| Column | Type | Description | Foreign Key | Required |
|---|---|---|---|---|
| id | serial | A unique identifier for each geographic location | | x |
| address_1 | text | Typically used for street address | | |
| address_2 | text | Typically used for additional detail such as building, suite, floor, etc. | | |
| city | text | The city field as it appears in the source data | | |
| state | text | The state field as it appears in the source data | | |
| zip | text | The zip or postal code | | |
| county | text | The county, if available | | |
| census_tract | text | The census tract if available | | |
| hsa | text | The Health Service Area, if available (originally defined by the National Center for Health Statistics) | | |
| country | text | The country if necessary | | |

- Used to store location information for patients, practitioners, and facilities
- One record for each geographic location in the data

**Table 16: Facilities**

| Column | Type | Description | Foreign Key | Required |
|---|---|---|---|---|
| id | serial | A unique identifier for each facility | | x |
| facility_name | text | Facility name, if available | | |
| primary_identifier | text | Primary facility identifier | | x |
| primary_identifier_type | text | Type of identifier specified in primary identifier field (UPIN, NPI, etc) | | x |
| secondary_identifier | text | Secondary facility identifier (Optional) | | |
| secondary_identifier_type | text | Type of identifier specified in secondary identifier field (UPIN, NPI, etc) | | |
| facility_type_concept_id | bigint | FK reference to concepts table representing the facility type | concepts | |
| specialty_concept_id | bigint | A foreign key to an identifier in the concepts table for specialty | concepts | |
| address_id | bigint | A foreign key to the address of the location of the facility | addresses | |

- Unique records for all the facilities in the data
- facility_type_concept_id should be used to describe the whole facility (e.g., Academic Medical Center or Community Medical Center). Specific departments in the facility should be entered in the contexts table using the care_site_type_concept_id field.

**Table 17: Concepts**

| Column | Type | Description | Foreign Key | Required |
|---|---|---|---|---|
| id | serial | Surrogate key for record (this is the concept_id) | | x |
| vocabulary_id | text | Unique text-string identifier of the vocabulary (see OMOP or UMLS) | vocabularies | x |
| concept_code | text | Actual code as text string from the source vocabulary (e.g., "410.00" for ICD-9) | | x |
| concept_text | text | Text descriptor associated with the concept_code | | x |

- Adapted from OMOP concept table (could add other fields, like domain, if needed)
- Can be created *de novo* for each data source or could use a different source like the National Library of Medicine Metathesaurus
- The mappings table can be used to establish relationships among concept ids

**Table 18: Vocabularies**

| Column | Type | Description | Foreign Key | Required |
|---|---|---|---|---|
| id | text | Short name of the vocabulary which acts as a natural key for record | | x |
| omopv4_vocabulary_id | int | Old ID used in OMOPv4 | | x |
| vocabulary_name | text | Full name of the vocabulary | | x |

- A list of vocabularies, currently adapted from the OMOP vocabulary table (e.g., ICD9)

**Table 19: Mappings**

| Column | Type | Description | Foreign Key | Required |
|---|---|---|---|---|
| concept_id_1 | bigint | FK reference to concepts table for the source concept | concepts | x |
| relationship_id | text | The type or nature of the relationship (e.g., "is_a") | | x |
| concept_id_2 | bigint | FK reference to concepts table for the destination concept | concepts | x |

- A set of relationships, currently adapted from the OMOP concept_relationship table
- This can be used to establish relationships between database-specific information and standardized information.
- It is preferable to store the raw data in the concepts table and establish a mapping to standard concepts in this table.
    - For example, if sex were coded as "male" and "female", these terms would be stored in the concepts table, and would be linked to standard concepts (if any) in the mappings table
    - This moves such "hidden" mappings from the ETL process to the data itself, makes ETL easier, and increases transparency and reproducibility of studies
